# *In vitro* and *in silico* characterization of competitive inhibition and repression of DUX4 target gene activation as a therapeutic approach for facioscapulohumeral muscular dystrophy (FSHD)

**DOI:** 10.64898/2026.08.04.742607

**Authors:** Heloise M Hoffmann, Alice Finkelstein, Amanuel Geremew, Katherine Xu, Vanessa Chiprez Meza, Ayushi Mohanty, Maria Fernanda Velásquez, Michael Liu, Alex Engel, Phillip Kyriakakis

## Abstract

Facioscapulohumeral muscular dystrophy (FSHD) is a rare neuromuscular disease caused by aberrant re-expression of the embryonic transcription factor DUX4 in skeletal muscle, which activates a toxic transcriptional program that drives progressive muscle wasting. No approved disease-modifying therapies currently exist. Prior work in mammalian and zebrafish models has shown that a truncated form of DUX4 retaining only its DNA-binding domain (DBD) lacks transactivation capacity and can suppress DUX4-FL-driven pathology; separately, dCas9/KRAB-based epigenetic repressors have demonstrated efficacy in silencing DUX4 expression, though CRISPR-based strategies face challenges from the repetitive nature of the D4Z4 locus, the immunogenicity associated with bacterial Cas proteins, and the payload limitations of gene delivery vehicles. Building on these findings, we corroborate that the DUX4 DBD, comprising both homeodomains, acts as a non-toxic competitive inhibitor of full-length DUX4 (DUX4-FL) at its genomic target sites, and extend this strategy by fusing the DBD to a human KRAB(ZNF10) domain, converting DUX4 from a transcriptional activator into a fully humanized epigenetic silencer of its own targets. Using a fluorescent DUX4-responsive reporter, we show that DBD alone produces dose-dependent repression of DUX4-FL transcriptional activity in HEK293T cells (200-fold at the highest inducible dose tested), while a constitutively expressed DBD-KRAB fusion produces significantly greater repression than DBD alone (949-fold versus 17-fold at a 25x molar ratio), with a similar trend observed in C2C12 myoblasts (47-fold versus 3.3-fold knockdown). To contextualize these findings and explore dosing considerations, we developed three complementary computational models – a transcription factor competitive binding model, a myotube diffusion model, and an ordinary differential equation (ODE) compartmental model – that illustrate how DBD concentration, intracellular diffusion, and population-level cell state transitions may relate to therapeutic efficacy. Together, these results corroborate and extend existing approaches into a single, fully humanized construct that may help circumvent the immunogenicity and delivery limitations of Cas-based systems.

## 1. INTRODUCTION

### 1.1 Facioscapulohumeral muscular dystrophy (FSHD)

Facioscapulohumeral muscular dystrophy (FSHD) is a rare, debilitating neuromuscular disorder affecting 4-12 in 100,000 individuals worldwide. The disease is characterized by progressive weakening of the muscles of the arms, shoulders, and face, but it can progress to any skeletal muscle, including the legs, trunk, and chest.^1–3^ FSHD often leads to significant disability, with about one in four patients requiring a wheelchair for mobility at some point during their lifetime and a majority of patients relying on some form of adaptive device, including leg and spinal braces, canes, walkers, and home modifications.^2,4^

Symptoms typically emerge in the second or third decade of life, but some patients experience early-onset FSHD, with symptoms beginning in the first decade and marked by severe, rapid progression and complications such as respiratory, visual, and auditory impairments.^2,5^ Despite major research efforts, there are currently no approved disease-modifying therapies for FSHD. The current standard of care relies on loose guidelines for physical therapy and symptom management.^1,3,4,6^ Here, we advance a strategy for repressing the toxic transcriptional network underlying FSHD.

### 1.2 DUX4 as the central driver of FSHD pathology

FSHD results from epigenetic dysregulation of the D4Z4 macrosatellite repeat array on chromosome 4, leading to aberrant re-expression of the embryonic DUX4 transcription factor in adult skeletal muscle.^2,7,8^ In people with FSHD, sporadic re-expression of DUX4 triggers toxic activation of downstream genes, initiating a cascade of pathogenic events including: activation of germline and early embryonic gene programs;^9,10^ oxidative stress and DNA damage;^11,12^ inhibition of myogenic differentiation;^13,14^ induction of inflammatory pathways;^15^ and progressive myocyte death.^16,17^ Even very low levels of DUX4 expression are sufficient to cause toxicity due to the downstream gene activation cascade, making it a challenging therapeutic target.^14^

Given its central role in disease, strategies targeting DUX4 are promising. However, existing strategies – including small molecules,^18^ RNA-targeting,^19–22^ or CRISPR-based approaches – face major limitations with durability, specificity, delivery, and immunological response,^21,23–26^ pointing to an unmet need for additional approaches that can safely and durably inhibit DUX4 activity in muscle in a clinically compatible manner.^6,27^

The toxic DUX4 transcription factor is composed of two functional domains:^13,28^ (1) an N-terminal region containing the DNA-binding domain (DBD), composed of two homeodomains, that targets specific genomic loci, and (2) a C-terminal transactivation domain (TAD) that recruits transcriptional machinery. The two are linked by an intrinsically disordered region (IDR). As previous work has shown, the toxic full-length DUX4 protein requires all domains intact (DBD, linker, and TAD) to perform its pathogenic function, and removing or neutralizing a single domain eliminates pathogenic activity.^13,28,29^

### 1.3 Harnessing DNA-binding specificity of DUX4 as an inhibitor of FSHD pathogenesis

Expression of a “short form” of DUX4 (DUX4-s) consisting only of the two DNA-binding homeodomains has been shown to suppress the pathological phenotype of full-length DUX4 misexpression in a zebrafish model of FSHD.^28^ As expected, DUX4-s lacked the ability to act as a transcriptional activator without the TAD. Additionally, previous work has demonstrated that truncated forms of DUX4 that retain only the N-terminal DBD are not toxic to myocytes.^28,30,31^ These findings motivated the idea to harness the DBD as a competitive inhibitor of DUX4 DNA-binding activity as a therapeutic candidate.

Hence, our strategy aims to block transcriptional network activation associated with FSHD via dominant-negative competition, using the DUX4 DBD, comprising both homeodomains, as a competitive inhibitor that binds to the same genomic loci as the toxic DUX4 protein and inhibits transcription. In addition, our strategy aims to further repress the activation of DUX4 targets via epigenetic repression at DUX4 target sites. The Krüppel-associated box (KRAB) is a powerful transcriptional repression domain that operates by complexing with the scaffolding protein KAP1, forming the scaffold for chromatin-modifying machinery that represses the transcription of downstream regions.^32–34^ KRAB-mediated repression has been used in the context of FSHD; in one study, a dCas9/KRAB transcriptional inhibitor complex targeted to the repetitive D4Z4 satellite array showed high efficiency in repressing DUX4 expression in FSHD myocytes.^35^ However, delivery of such a large system and the immunogenic effects of the bacterial protein Cas9 could be challenging from a clinical standpoint.

Here, we present a fully humanized, compact epigenetic repressor of the DUX4 transcriptional network by fusing the DBD (hereby referenced as “DBD”) to a KRAB(ZNF10) module, linked via a truncated form of the intrinsically disordered region (IDR) that typically links the DBD and TAD (DBD-KRAB). We have converted DUX4 from a transcriptional activator to a repressor that silences rather than upregulates its own targets. Our dual approach is expected to effectively block the pathogenic pathway regulated by DUX4 by (a) inhibiting DUX4 binding to target DNA sites via competitive inhibition, and (b) converting the chromatin architecture of DUX4 target genes to a repressive state.

A key advantage of this approach over genome and epigenome-editing strategies such as CRISPR-based systems is its ability to antagonize pathogenic DUX4-FL activity at all of its target binding sites. CRISPR-based interventions are constrained to correcting a single, predefined sequence, and the highly repetitive region at the D4Z4 locus is difficult to target.^36^ While CRISPR-based interventions can target unique flanking sequences of the D4Z4 locus, such as the terminal exon distal to the 4q35 array,^37^ nuclease-based approaches carry risks of off-target DNA cleavage, and non-cleaving epigenetic or base-editing variants remain limited by the restrictive heterochromatic architecture of the D4Z4 locus.^38^ Furthermore, recent Telomere-to-Telomere (T2T) genomic mapping has revealed that D4Z4-like elements and highly homologous fragments of the terminal exon are scattered across at least ten additional chromosomes,^39^ compounding the risk of unpredicted off-target guide RNA binding.

The approach using DBD and DBD-KRAB as competitive inhibitors of DUX4-FL avoids this challenge, as DBD and DBD-KRAB are expected to recapitulate the binding specificity of DUX4-FL across its genomic targets with near-equivalent affinity without requiring patient-specific sequence optimization. Additionally, the bacterial origin of Cas proteins is associated with documented immune responses,^40–43^ and preexisting adaptive immunity in a substantial proportion of the human population,^44,45^ both of which risk compounding the pathological effects of DUX4-FL overexpression. Although strategies to mitigate these immunological barriers are under investigation,^46^ the use of endogenous human-derived proteins like DBD and DBD-KRAB circumvents these issues, thereby paving the way for gene therapies for FSHD potentially associated with high specificity and low immunogenic concerns.

## 2. METHODS

### 2.1 Plasmids & Cloning

DNA plasmids were constructed via Gibson Assembly using NEBuilder HiFi DNA Assembly Master Mix (New England Biosciences). All transformations were carried out in DH5α or DH10□ cells.

Plasmid pSI-007 (a constitutive DUX4-FL construct with DUX4-FL driven from the SV40 promoter, referred to as DUX4-FL) consists of an mTAG-BFP2 sequence followed by a P2A linker upstream of the DUX4-FL sequence. The DUX4-FL sequence and backbone of pSI-007 were amplified from pCW57.1-DUX4-CA (Addgene plasmid #99281),^47^ and the TRE promoter in pCW57.1-DUX-CA was replaced by a constitutive SV40 promoter PCR-amplified fragment derived from pPK101.^48^

Plasmid pSI-008 (a doxycycline-inducible DBD construct, iDBD) was constructed from pCW57.1-DUX4-CA by inserting an mNeonGreen sequence followed by a peptide 2A (P2A) linker upstream of the DBD sequence. The DBD sequence consists of the first 217 N-terminus amino acids of DUX4-FL and excludes its transcriptional activation domain (TAD).^28^

Plasmid pSI-017 (a constitutive DBD construct with DBD driven from the SV40 promoter, cDBD) was constructed by amplifying pSI-008 without the TRE promoter and assembling the resulting fragment with an SV40 promoter PCR-amplified fragment derived from pSI-007.

Plasmid pSI-009 (DUX4 fluorescent reporter) was constructed from pGL4-6X-DUX4-ffy, a gift from Michael Kyba,^29^ which contains 6 repeated DUX4-binding sites upstream of a minimal promoter and a pLuc sequence encoding luciferase. To make pSI-009, the backbone of pGL4-6X-DUX4-ffy, excluding the fLuc sequence, was amplified and assembled with mScarlet (from plasmid pPK381^49^) in place of fLuc.

To generate plasmid pSI-019 (DUX4-KRAB), the Krüppel Associated Box (KRAB) repressor domain was appended to the end of the DBD sequence from pSI-017. The KRAB sequence was synthesized by IDT based on plasmid pHAGE EF1α dCas9-KRAB (Addgene plasmid #50917). The KRAB fragment was then assembled with the amplified backbone of plasmid pSI-017.

#### Plasmid Availability

All plasmids generated in this study have been deposited to GenBank under the following accession numbers: pSI-007 SV40-mTag BFP2-DUX4-FL (PZ794249); pSI-008 tet-mNG-P2A-DBD (PZ794250); pSI-009 6X-DUX4-UAS-mScarlet (PZ794251); pSI-017 SV40-mNG-P2A-DBD (PZ794252); pSI-019 SV40-mNG-P2A-DBD-ZNF10 KRAB (PZ794253); pPK-205 20XUAS_luc (PZ794254).

### 2.2 Cell Culture

Human embryonic kidney (HEK293T) cells and mouse myoblast (C2C12) cells (a gift from Helen Blau) were maintained under standard culture conditions (5% CO2 and 37ºC) with Dulbecco’s Modified Eagle’s Medium (DMEM; Thermo Fisher Scientific) supplemented with 10% Fetal Bovine Serum (FBS; Thermo Fisher Scientific) and 1% Penicillin-Streptomycin (Thermo Fisher Scientific).

### 2.3 Competitive inhibition of DUX4 reporter activity

Competitive inhibition experiments in HEK293T cells were conducted by transfection in biological replicates. 5 x 10^4^ HEK293T cells were plated per well 24 hours before transfection. Lipofectamine 3000 (Thermo Fisher Scientific) was used according to the manufacturer’s protocol.

For transfections with DBD under inducible control (**Fig. 1C, D**), pSI-007 (34.2 ng), pSI-009 (20 ng), and pSI-008 at varying construct:DUX4-FL molar ratios (1:1, 10:1, and 25:1) were co-transfected in HEK293T cells. Filler DNA (pPK-205) was used to normalize the total mass of DNA transfected. Media was replaced six hours after transfection with media containing doxycycline (250 ng/mL) to induce expression of iDBD.

**Figure 1.**
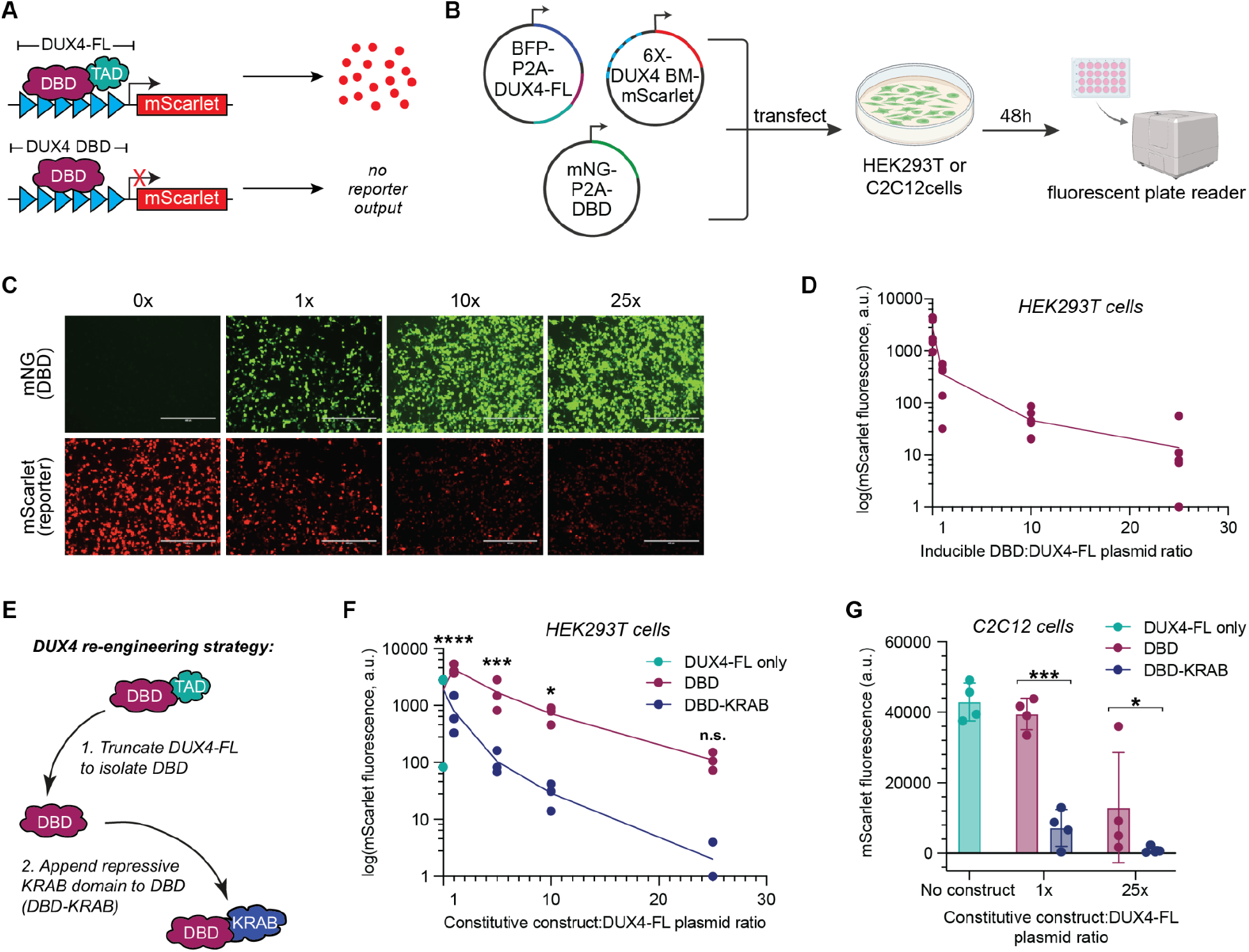
DBD and DBD-KRAB repress DUX4-FL-driven reporter activity in a dose-dependent manner in HEK293T and a skeletal muscle cell model. **(A)** Schematic of the DUX4-FL reporter system. Reporter features 6X DUX4 DNA binding motifs (*blue triangles*) upstream of mScarlet. **(B)** Diagram representing the method to assess reporter inhibition: plasmids encoding constitutive DUX4-FL, the mScarlet reporter, and the DBD construct were co-transfected into HEK293T cells. The molar ratio of DBD:DUX4-FL was varied with constant DBD induction levels. **(C)** Reporter (*red*) and DBD construct (*green*) expression was observed in HEK293T cells via fluorescence microscopy. *Scale bar = 400 μm*. **(D)** Reporter activity measured in HEK293T cells 48h after transfection. The line connects means at each dose. **(E)** DBD-KRAB construct was engineered by appending a ZNF10 KRAB domain to the C-terminus of the DBD. **(F)** Reporter activity was measured in HEK293T cells co-transfected with plasmids encoding constitutive DUX4-FL, mScarlet reporter, and constitutive therapeutic constructs (DBD or DBD-KRAB), at varying construct:DUX4-FL molar ratios. The lines connect the means at each dose. A two-way ANOVA was used to analyze the results; stars represent a significant differences between DBD and DBD-KRAB treatment at each dose (*\*p<0*.*05, ***p<0*.*001, ****p<0*.*0001*). **(G)** Reporter activity was measured in C2C12 cells co-transfected with reporter, DUX4-FL, and DBD constructs. Error bars = standard deviation. A two-way ANOVA was run; stars represent a significant difference between DBD and DBD-KRAB treatment at each dose (*\*p<0*.*05, ***p<0*.*001*).

For transfections with constitutive DBD and DBD-KRAB (**Fig. 1F**), pSI-007 (34.2 ng), pSI-009 (20 ng), and pSI-017 or pSI-019 at varying cDBD/cDBD-KRAB:DUX4-FL molar ratios (1:1, 5:1, 10:1, and 25:1) were co-transfected in HEK293T cells. pPK-205 was used to normalize the total mass of DNA transfected.

To quantify reporter repression, 48 hours after transfection, mScarlet (reporter) and GFP (iDBD, cDBD, or DBD-KRAB) expression was quantified using an area scan on a Synergy H1 microplate reader (Agilent).

For competitive inhibition experiments in C2C12 cells, reverse transfections were conducted in 24-well plates in biological replicates. At the time of transfection, 20,000 C2C12 cells were plated in 1mL of DMEM. Lipofectamine 3000 (Thermo Fisher Scientific) was used for transfection with plasmids encoding DUX4-FL (pSI-007, 29.74 ng), the mScarlet reporter (pSI-009, 32.66 ng), and cDBD (pSI017) or DBD-KRAB (pSI-019) constructs at varying construct:DUX4-FL molar ratios (1:1, 5:1, 10:1, and 25:1). pPK-205 was used to normalize total mass of DNA transfected.

To quantify reporter repression, 48 hours after reverse transfection, after cells were dense and many myotubes had formed, the media was aspirated and washed with 1X PBS buffer. Cells were lysed with 20ul of 1X Passive Lysis Buffer (Promega Corp) and frozen at -20°C. The lysed cell solution was thawed between 4 and 76 hours after freezing, and suspensions were transferred to a 384 square well plate (Greiner Bio-One) and assayed using a Tecan Infinite 200 PRO M Plex plate reader.

### 2.4 Statistical analysis and data visualization of DUX4 reporter assay data

Plate reader data for experiments in HEK293T and C2C12 cells were normalized for each replicate as follows: Background subtraction was completed by subtracting fluorescence in the mScarlet channel in the baseline reporter-only control from the mScarlet fluorescence in each experimental condition, thereby accounting for leaky reporter expression and background fluorescence. The final value represents the normalized mScarlet expression (a.u.) in each condition.

For normalized fluorescence values below zero after background subtraction, the =MAX(value,1) function was used to convert the lowest possible fluorescence readout to 1, since a negative a.u. value indicates fluorescence close to background levels.

To determine the average fold-knockdown of reporter expression in each experiment, the average normalized mScarlet expression value for the DUX4-FL-only condition was divided by the average normalized mScarlet expression value for the highest construct:DUX4-FL ratio tested (25x). Two-way ANOVA was performed to determine whether there were statistically significant differences in reporter expression between doses and between treatments. Tukey’s multiple comparisons test was used to determine which data points were significantly different.

Data were visualized using GraphPad Prism, version 11.0.0 for Mac, GraphPad Software, 2026. Figures were produced using Adobe Illustrator, version 30.3 for Mac, Adobe Inc., 2026.

### 2.5 Transcription Factor Competitive Binding Model

#### Single-Site Binding Model

Competitive occupancy at a single DUX4 consensus site was modeled using the equilibrium binding equation derived from mass-action kinetics:^50^

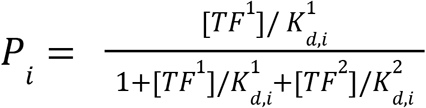

In our case, [TF^1^] represents DUX4-FL concentration. [TF^2^] represents DBD concentration. K_D_ is the dissociation constant, assumed to be the same for both DBD and DUX4-FL, and is 1.4 μM.^51^ DUX4-FL concentrations were modeled across three biologically meaningful regimes:

- High: [FL] = 1 × 10⁻^3^ M (overexpression conditions of transient transfection; [FL] ≫ K_D_)
- Mid: [FL] = 1 × 10⁻^6^ M ([FL] ≈ K_D_; approximating low micromolar physiological expression)
- Low: [FL] = 1 × 10⁻^9^ M ([FL] ≪ K_D_; approximating nanomolar physiological expression) DBD concentrations were varied up to 1,000-fold above [FL] to map the full inhibition landscape across each regime.

### 2.6 Myotube Diffusion Model

The molecular weight of DBD was computed from its amino acid sequence using Biopython’s `ProteinAnalysis` module and compared against well-characterized fluorescent proteins, mCherry, tdTomato, and DsRed, for which intracellular diffusion coefficients in muscle fibers have been experimentally determined.^52^ The DBD diffusion coefficient was estimated via linear interpolation between the mCherry and tdTomato values based on molecular weight.

Using this estimated coefficient, we numerically solved the one-dimensional diffusion equation along a model muscle fiber, tracking the DBD concentration profile at multiple time points. We additionally computed steady-state concentration profiles using an exponential decay approximation for DBD and all three reference proteins. All simulations were implemented in Python using Biopython and Matplotlib.

### 2.7 Ordinary Differential Equation (ODE) Compartmental Model

An ODE-based compartmental model was developed to simulate the dynamic evolution of DUX4-FL and DBD interactions across a population of cells over a 48-hour period.^53^ Five discrete cell states were defined: Susceptible (S, no DUX4-FL expression), Exposed (E, DUX4-FL mRNA present but no detectable protein), Infected (I, DUX4-FL mRNA and protein present), Recovered (R, declining DUX4-FL protein), and Dead (D, cell death due to DUX4-FL cytotoxicity).

The DBD + FL condition was modeled within the FL-positive compartments rather than as a separate sixth cell state. Specifically, cells in the Exposed or Infected compartments could contain both DUX4-FL and DBD, and the effect of DBD was represented by reducing the effective DUX4-FL activity according to the DBD:FL ratio derived from the competitive binding model. Thus, DBD does not change whether a cell is classified as FL-positive; instead, it changes the probability that FL-positive cells progress toward cytotoxicity.

State transition rates were governed by a system of ODEs adapted from the in silico FSHD muscle fiber model of Cowley et al.^53^ Compared with the original framework, we introduced a DBD-dependent modulation of the infection/progression rate, β_eff, such that increasing DBD relative to FL reduced the effective rate of progression from FL expression toward downstream toxic states. This allowed the model to simulate untreated FL-only conditions, DBD-only conditions, and combined DBD + FL conditions within the same compartmental structure. **Fig. 2D** shows the untreated baseline trajectory under this fitted, DBD-independent infection rate. Explicit coupling of β to the DBD:FL ratio is a planned extension of this framework; the baseline model and fitting procedure described here are available in the linked repository.

**Figure 2.**
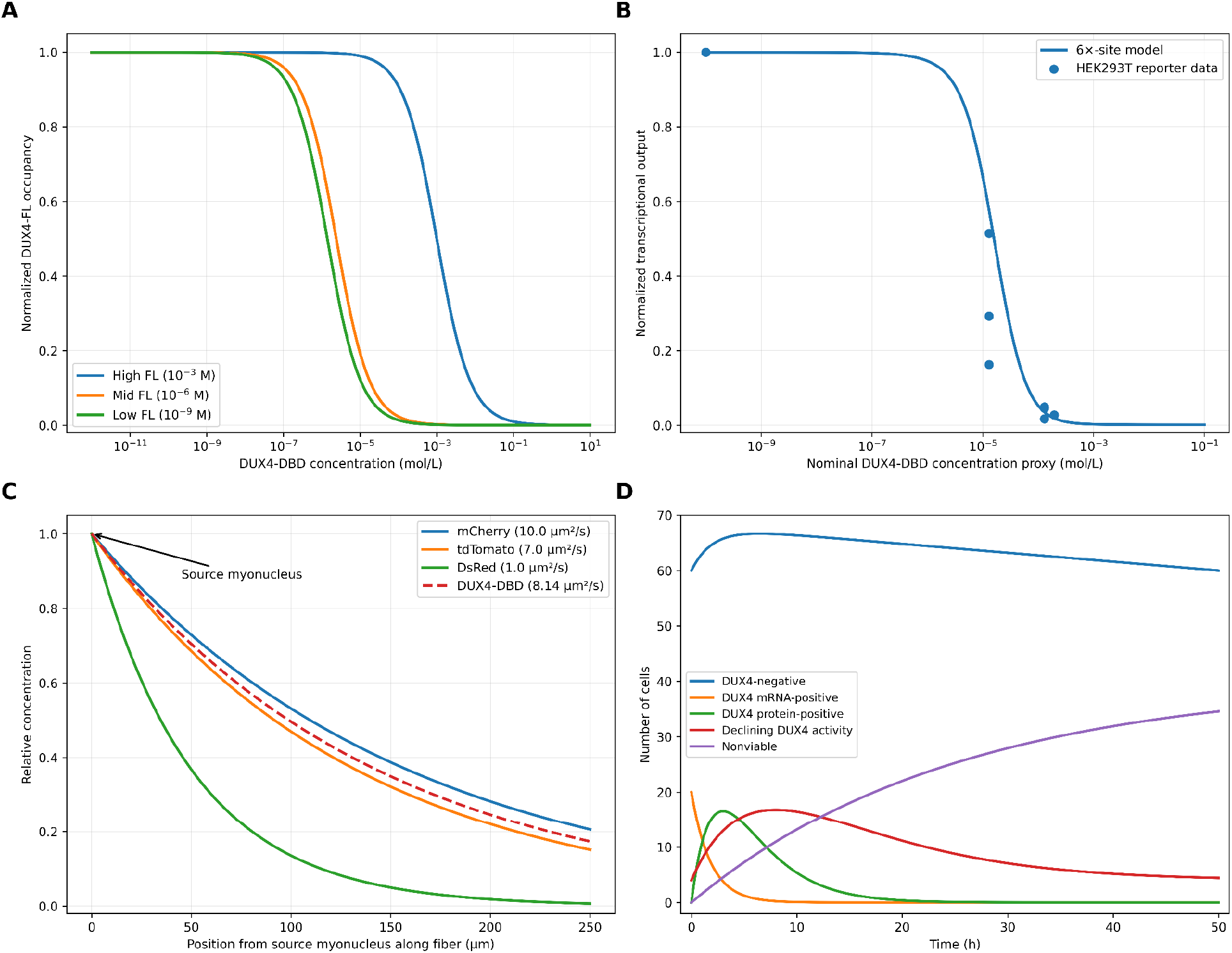
Computational modeling predicts concentration-, spatial-, and time-dependent DBD inhibition of DUX4-FL activity. **(A)** Competitive-binding model predicts normalized DUX4-FL-driven transcriptional output as a function of DBD concentration for a fixed 2X DUX4-binding-site promoter under high, mid, and low DUX4-FL regimes. Each curve was normalized to its own no-DBD condition after subtraction of the modeled basal output. **(B)** Comparison of the 6× DUX4-binding-site competitive-binding model with normalized reporter measurements from three independent HEK293T transient-transfection experiments. Experimental x-axis positions are based on nominal DBD:FL plasmid molar ratios and should not be interpreted as measured intracellular concentrations. **(C)** Illustrative steady-state concentration profiles generated using a one-dimensional diffusion-with-loss model for DBD and fluorescent reference proteins. Larger diffusion coefficients produce broader spatial profiles under the shared loss-rate assumption. The estimated diffusion coefficient of DBD was obtained by molecular-weight interpolation and should be interpreted as a sensitivity-analysis parameter rather than a direct measurement. **(D)** Exploratory baseline ODE simulation of transitions among DUX4-FL-negative, DUX4-FL mRNA-positive, DUX4-FL protein-positive, declining DUX4-FL-activity, and nonviable states over 48 hours without an explicitly modeled DBD intervention.

To capture the inherent stochasticity of DUX4-FL transcriptional bursting, we incorporated stochastic gene expression using the Poisson-Beta distribution framework.^54^ Posterior parameters (α and β) were derived from a Poisson-Beta MCMC model fit to simulated mRNA transcript counts. Base DUX4-FL and DBD expression parameters were specified as model inputs rather than directly measured biological rates. Because the provenance of the original values could not be independently verified, they were treated as exploratory assumptions and varied in sensitivity analyses.^55^ The infection rate β—serving as a proxy for disease progression—was modulated dynamically by the real-time DBD:FL ratio, directly coupling the competitive binding dynamics from Section 2.2 into the compartmental framework.

#### Code Availability

All Python code used to implement the competitive binding model, diffusion simulations, ODE compartmental model, and Markov chain simulations is available on GitHub at: https://github.com/aksarkar/poisbeta

## 3. RESULTS

### 3.1 DBD inhibits DUX4-FL transcriptional activity in HEK293T cells

We first aimed to evaluate whether the DBD could repress DUX4-mediated transcriptional activation in a proof-of-concept system. To this end, we engineered a fluorescent DUX4 reporter (pSI-009) that expressed mScarlet under the control of six tandem DUX4 binding sites, constructed from a previously validated DUX4-sensitive reporter.^1^ We first set out to test whether a truncated DBD construct (including only the 1-217 N-terminus amino acids (AA)), lacking the transcriptional activation domain, could competitively inhibit full-length DUX4-FL. Therefore, we transfected HEK293T cells with the mScarlet reporter, DUX4-FL, and an mNeonGreen (mNG)-DBD construct (DBD-P2A-mNG) at increasing DBD:DUX4-FL molar ratios (**Fig. 1A, B**). mNeonGreen fluorescence, representing DBD construct expression, increased with increasing DBD:DUX4-FL molar ratios, confirming dose-dependent expression of the DBD construct (**Fig. 1C**). Importantly, DUX4 reporter activity (measured by mScarlet fluorescence) progressively decreased as DBD:DUX4-FL molar ratio increased, indicative of competitive inhibition of DUX4-FL binding to target sites (**Fig. 1C-D**). Notably, at the highest inducible DBD concentration, a 200-fold decrease was observed compared to the condition with no therapeutic construct.

### 3.2 DBD-KRAB fusion protein enhances DUX4-FL repression

Because even low levels of DUX4 are sufficient to drive FSHD pathogenesis,^2^ we explored further improving the inhibitory effect of the DBD construct by fusing a ZNF10 KRAB transcriptional repressor module to the C-terminus of DBD, generating the DBD-KRAB construct (pSI-019) (**Fig. 1E**). For this experiment, we pivoted to co-transfection with inhibitory constructs under the same constitutive SV40 promoter as DUX4-FL plasmid in order to more precisely control the molar ratio of construct:DUX4-FL expression levels. In HEK293T cells, DBD-KRAB produced a significantly greater knockdown of mScarlet reporter activity than DBD (**Fig. 1F**). At a 25x DBD/Dux4FL or DBD-KRAB:DUX4-FL molar ratio, relative to DUX4-FL alone, DBD-KRAB caused a 949-fold decrease in reporter expression, compared to a 17-fold decrease with DBD treatment. There was a significant difference in reporter repression mediated by DBD versus DBD-KRAB at 1x, 5x, and 10x doses (*p<0*.*0001, p<0*.*001, p<0*.*05*, respectively). At the highest dose tested (25x), DBD-KRAB treatment brought fluorescence to the background levels of untransfected conditions (**Fig. 1F**). However, this effect was not significantly different from the DUX4-FL-only condition; in fact, within each treatment condition, there was no significant difference between the DUX4-FL-only condition and each dose.

### 3.3 DBD-based constructs repress reporter in muscle-specific system

Finally, we asked whether our system could achieve efficacy in a skeletal muscle lineage cellular environment, which has known differences in the core transcriptional machinery.^57,58^ To test this, we transferred our reporter assay system into C12C12 cells, an immortalized murine myoblast model.^59^ C2C12 cells transfected with reporter, DUX4-FL, and therapeutic plasmid constructs showed inhibition of DUX4-mediated transactivation in the skeletal muscle environment, with the added repression from the KRAB module outperforming DBD alone. At the highest plasmid ratio of construct:DUX4-FL tested, DBD caused, on average, a 3.3-fold knockdown of reporter expression relative to the condition with no therapeutic construct, while DBD-KRAB caused a 47-fold knockdown (**Fig. 1G**). There was a significant difference in DBD vs DBD-KRAB -mediated reporter repression at both doses tested in this experiment (1x and 25x; *p<0*.*001, p<0*.*05*, respectively). Additionally, within the DBD treatment condition, there was a significant difference from the DUX4-FL-only condition for the condition treated with 25x construct (*p<0*.*0001*), and within the DBD-KRAB treatment condition, both the 1x and 25x constructs were statistically significantly different from the DUX4-FL-only condition (*p<0*.*0001* in both cases).

### 3.4 Mathematical model of transcription factor competitive binding predicts concentration-dependent inhibition

Translating DBD competitive inhibition into a viable therapeutic strategy requires a sufficient quantity of DBD to block expression of DUX4-FL target genes. Three principal challenges motivate a computational framework. <u>First</u>, DUX4-FL expression is highly variable between patients and across individual nuclei within a single fiber.^15^ <u>Second</u>, skeletal muscle fibers are syncytial, and DUX4-FL expression is rare, sporadic, and restricted to only a small fraction of myonuclei at any given time.^8^ <u>Third</u>, therapeutic AAV-based gene delivery is expected to transduce only a subset of myonuclei, while nuclear-targeted proteins can propagate across myonuclear domains within a shared muscle fiber, creating spatial heterogeneity in whether DBD reaches the nuclei where DUX4-FL is active.^60^ To address these challenges in an integrated manner, we developed three complementary models: (i) a transcription factor competitive binding model, (ii) a myotube protein diffusion model, and (iii) an ordinary differential equation (ODE) compartmental model. Together, these models are designed to answer the central question: what concentration of DBD is required to achieve a defined inhibition threshold of DUX4-FL-driven transcription, and how does this threshold vary with patient-specific expression levels and cellular context?

First, the transcription factor binding model quantified the DBD threshold required for inhibition across various baseline DUX4-FL concentrations (**Fig. 2A**). For this sensitivity analysis, the model represents a simplified promoter containing two identical DUX4 target sequences, rather than the variable motif number, spacing, orientation, and sequence context of endogenous DUX4-regulated genes. The DUX4-FL values were chosen as sensitivity-analysis regimes relative to K_D_, rather than as directly measured physiological concentrations. Under the high-FL model regime, DUX4-FL concentration was fixed at 10⁻^3^ M, several orders of magnitude above the assumed dissociation constant for DUX4 DNA binding, K_D_ ≈ 1.4 µM.^51^ In this high-FL setting, near-maximal transcriptional output was maintained until DBD concentrations reached the 10⁻^5^-10⁻^3^ M range, reflecting the mass-action advantage DUX4-FL has when present in large excess. At the mid-FL regime (10⁻^6^ M), approximating K_D_, the inhibition curve shifted leftward, with suppression beginning at lower DBD concentrations. At the low-FL regime, 10⁻^9^ M, lower absolute DBD concentrations were sufficient to shift binding toward DBD-dominant occupancy.

Across the three modeled DUX4-FL regimes, increasing baseline DUX4-FL shifted the inhibition curve toward higher required DBD concentrations (**Fig. 2A**). Thus, the model predicts that the DBD level required for a given degree of relative suppression depends strongly on the starting DUX4-FL burden.

### 3.5 Experimental validation against HEK293T reporter data

To evaluate whether the binding model captured the competitive dynamics of DUX4 inhibition, we compared model-predicted transcriptional output with normalized mScarlet reporter signal from three independent HEK293T transient-transfection experiments (**Fig. 2B**). DBD (pSI-008) was expressed from a doxycycline-inducible TRE promoter, induced at [25 ng/µL] doxycycline six hours post-transfection, in all conditions, transfected at plasmid molar ratios of 1:1, 5:1, 10:1, and 25:1 relative to the DUX4-FL plasmid. Because absolute intracellular DBD concentrations were not directly measured, experimental data points were overlaid onto the model curve by treating the nominal DBD:FL molar ratio as a proxy for relative DBD concentration, an approximation that assumes proportional plasmid uptake and expression across conditions.

Using the concentration-axis calibration described above, the 6X binding-site model showed approximate agreement with the HEK293T reporter data (**Fig. 2B**). This calibration assumed that equal molar amounts of the DUX4-FL and DBD plasmids produced approximately proportional intracellular protein concentrations under the experimental expression conditions. Under this assumption, the observed inhibition trend was consistent with increasing DBD:FL molar ratios, with near-complete suppression at the highest tested ratio. Because intracellular protein concentrations were not directly measured, the data do not establish absolute DUX4-FL or DBD concentrations, including for the transient-transfection conditions used here.

The model reproduced the general concentration-dependent inhibition trend observed in the HEK293T reporter experiments (**Fig. 2B**), particularly the transition from partial inhibition at intermediate DBD:FL ratios to near-complete suppression at the highest tested ratios. However, the experimental points did not follow the model curve exactly, especially at intermediate ratios, indicating that the simplified equilibrium model did not fully capture the observed inhibition response

### 3.6 Complementary models extend the framework to space and time

The binding model, while mechanistically grounded, captures only a static equilibrium view of competitive inhibition in an isolated cell/compartment. Two complementary models address the dimensions the first model necessarily omits.

The myotube diffusion model addresses the spatial challenge inherent to syncytial muscle fibers. Although the fraction of myonuclei containing transcriptionally active AAV episomes within an individual targeted muscle fiber has not been well quantified and likely varies with vector, dose, delivery route, and tissue context, DBD protein produced by episome-containing myonuclei may need to diffuse through the shared cytoplasm to reach nuclei where FL is actively bursting - that is, undergoing stochastic pulses of transcriptional activity characteristic of DUX4-FL expression in FSHD myonuclei.^8^

The molecular weight of DBD was computed as 45.04 kDa using Biopython’s ProteinAnalysis module, placing it between mCherry (28.8 kDa) and tdTomato (55.0 kDa) and below dsRed (which forms a 110kDa tetramer), fluorescent proteins whose intracellular diffusion coefficients in muscle fibers have been experimentally characterized.^52^ For the purposes of the diffusion model, each episome-containing myonucleus was treated as a localized source of newly synthesized DBD protein. This assumes that transgene transcription occurs within the transduced myonucleus and that at least a portion of translation occurs within its surrounding myonuclear domain before the protein disperses through the shared cytoplasm. This is a simplifying assumption, as skeletal-muscle mRNAs can undergo microtubule-dependent transport away from their source nuclei, and the localization of AAV-derived DBD transcripts has not been directly measured.^61^ The DBD diffusion coefficient was estimated via linear interpolation between these two reference proteins by molecular weight, yielding an estimated coefficient of 8.14 µm^2^/s. This interpolation assumes that DBD cytoplasmic diffusion scales with molecular weight similarly to these fluorescent reference proteins. DBD stability, binding interactions, and intracellular transport were not independently characterized. Numerical simulation of the one-dimensional diffusion equation along a 250 µm model myotube was used as a first-order approximation of longitudinal DBD transport (**Fig. 2C**). This simplification is motivated by the elongated geometry of myotubes and focuses on diffusion between myonuclear domains along the fiber axis, but it does not capture radial concentration gradients, variation in fiber diameter, branching, or obstruction by intracellular structures. Under these assumptions, the model predicted that DBD could spread from a localized production source over biologically relevant distances and timescales.

Finally, the ODE compartmental model illustrates the time-dependent behavior of the modeled cellular states in the absence of an explicit DBD intervention (**Fig. 2D**). Cells transition between Susceptible (DUX4-FL-negative), Exposed (DUX4-FL mRNA-positive), Infected (DUX4-FL protein-positive), Recovered (declining DUX4-FL activity), and Dead (nonviable) states over 48 hours under the selected baseline parameters.

## 4. DISCUSSION

The goal of this study was to establish a novel method for the treatment of FSHD, harnessing the native DUX4 DNA-binding domain (DBD) as a competitive inhibitor of the pathogenic DUX4-FL transcription factor at the root of the disease. Our results corroborate prior work in mammalian and zebrafish models demonstrating that DBD acts as a suppressor of DUX4-FL-mediated transcriptional activation.^9,13,30,31^ By appending a repressive Krüppel-associated box (KRAB) domain to the DBD, we have enhanced its inhibitory potential, leading to significantly elevated reporter repression compared to DBD alone.

All cell-based experiments used transient transfection and yielded DUX4-FL concentrations that exceeded physiological FSHD levels; therefore, further validation is necessary to advance the translational potential of this work. Validation in human FSHD-patient-derived myotubes, which endogenously express DUX4-FL, is essential for characterizing the therapeutic potential of DBD and DBD-KRAB in a more physiologically relevant context. Further, using patient-derived myotubes can demonstrate the effects of DBD and DBD-KRAB in tissues from donors with different DUX4-FL expression levels. Transcriptional changes across the whole genome should be quantified, and the genome-wide binding specificity of the DBD and DBD-KRAB constructs should be profiled in order to identify potential off-target interactions.

Beyond the experimental validation, our computational framework provides additional insight into the dosing requirements and delivery constraints of this approach. Our computational binding model predicts how effective DBD concentrations are compared to baseline DUX4-FL levels. The assumption of equal dissociation constants for DBD and FL remains to be confirmed by direct measurement. Similarly, the absolute concentration scale used to align the binding model with the HEK293T reporter data was not directly measured and should be interpreted only as an order-of-magnitude curve-alignment assumption, rather than as an estimate of intracellular DUX4-FL or DBD concentration. The modeled low-FL result should likewise be interpreted relative to its assumed DUX4-FL baseline rather than as an independently defined therapeutic dose. More generally, because the high-, mid-, and low-FL values were selected as sensitivity-analysis conditions rather than measured physiological concentrations, the corresponding curves are intended to illustrate concentration dependence rather than define therapeutic dose targets. Discrepancies between model predictions and experimental observations likely reflect factors not captured by a static equilibrium framework, including differences in DBD and FL protein half-lives, non-specific nuclear interactions, and transcriptional bursting dynamics, and suggest that therapeutic dosing targets should lean toward the higher end of model predictions to account for biological noise.

The diffusion model supports the plausibility of DBD propagation between neighboring myonuclear domains, but they should be interpreted as a simplified estimate of longitudinal spreading rather than evidence that partial myonuclear transduction alone would provide complete therapeutic coverage. Specifically, this model predicts that even at 250µm from the transduced nucleus, a nucleus could receive ∼20% of the dose of the transduced nucleus, which is far enough to expose many nuclei to DBD or DBD-KRAB from a single transduced nucleus.^52^ However, it does not address fibers receiving no transduction at all. Because AAV-mediated skeletal-muscle transduction can vary with capsid, dose, administration route, and muscle context, some fibers or myonuclear regions may remain below an effective expression threshold.^62^ Diffusion efficiency alone cannot guarantee therapeutic coverage. Future work should characterize the minimum transduction frequency required for efficacy, both computationally and in patient-derived myotube systems, and/or by optimizing AAV capsids for *in vivo* delivery.

Finally, the current ODE simulation illustrates the temporal structure of the modeled DUX4-associated cellular states rather than quantifying the effect of DBD treatment. Incorporating a validated DBD:FL-dependent inhibition term and comparing trajectories across DBD concentrations will be necessary to determine how competitive inhibition alters population-level progression. More broadly, future population-level modeling should incorporate experimentally supported transition rates and cellular heterogeneity to better inform dosing and delivery strategies for *in vivo* studies. To model KRAB inhibition, where the chromatin is modified, making the DNA inaccessible to DUX4-FL, would be an interesting follow-on for understanding how it can influence disease progression compared to using DBD alone.

Together, these findings establish a foundation for a fully humanized therapeutic that represses the DUX4-FL-activated transcriptional program through competitive inhibition of DUX4-FL binding sites and epigenetic silencing, while potentially circumventing immunogenic concerns and gene delivery payload size constraints.

## 5. ACKNOWLEDGEMENTS

Our team is endlessly grateful for the support we have received from our mentors on this project. We thank Drew Endy, PhD for his oversight and support of this project. We also thank Samuel King and Cyrus Knudsen for their mentorship, and specifically their contributions to the dry lab components of this research. We thank Helen Blau, PhD for her priceless mentorship and the donation of the C2C12 cells. We appreciate the advisory support and technical feedback provided by previous iGEM members Julia Vu, Ngoc Tran, and Nick Murphy. We thank Peter Jones, PhD for his insights on the future directions of our work. Additionally, this work would not have been possible without the Uytengsu Teaching Lab at Stanford University, and in particular Mong Saetern and Jeffrey Tok, who provided lab space and invaluable support throughout this project. We thank Dr. Christopher Emig for his direction of the iGEM program at Stanford and his support for the publication of this work. We thank iGEM and TwistBioscience for their material contribution to DNA synthesis that supported this research. We also thank the Stanford Bioengineering department for providing funding for conference travel and reagents. Our team is grateful for the donation of the reporter plasmid pGL4-6X-DUX4-ffy from the Michael Kyba Lab of the University of Minnesota Twin Cities. Our team is deeply grateful for the contributions of the FSHD Society to our outreach work with patients and to the FSHD patient and caregiver community, whose resilience and hope inspire optimism.

## 6. AUTHOR CONTRIBUTIONS

Heloise Hoffmann (HH), Alice Finkelstein (AF), and Phillip Kyriakakis (PK) formulated the central research questions. HH, AF, PK, Vanessa Chiprez Meza (VC), and Ayushi Mohanty (AM) designed experiments. HH, AF, and PK interpreted data from cell experiments and performed troubleshooting. Katherine Xu (KX) and Maria Fernanda Velásquez (MV) planned and executed computational modeling and experiments. PK, AF, VC, and HH designed and constructed plasmids. HH, AF, VC, AM, Amanuel Geremew (AG), and Michael Liu (ML) performed cell experiments. HH, PK, KX, AF, and Alex Engel (AE) wrote the manuscript. AE and PK provided guidance and mentorship on experimental design and project development. All authors reviewed the manuscript and provided feedback.

